# Inhibitor Binding Modulates Protonation States in the Active Site of SARS-CoV-2 Main Protease

**DOI:** 10.1101/2021.01.12.426388

**Authors:** Daniel W. Kneller, Gwyndalyn Phillips, Kevin L. Weiss, Qiu Zhang, Leighton Coates, Andrey Kovalevsky

## Abstract

The main protease (3CL M^pro^) from SARS-CoV-2, the virus that causes COVID-19, is an essential enzyme for viral replication with no human counterpart, making it an attractive drug target. Although drugs have been developed to inhibit the proteases from HIV, hepatitis C and other viruses, no such therapeutic is available to inhibit the main protease of SARS-CoV-2. To directly observe the protonation states in SARS-CoV-2 M^pro^ and to elucidate their importance in inhibitor binding, we determined the structure of the enzyme in complex with the α-ketoamide inhibitor telaprevir using neutron protein crystallography at near-physiological temperature. We compared protonation states in the inhibitor complex with those determined for a ligand-free neutron structure of M^pro^. This comparison revealed that three active-site histidine residues (His41, His163 and His164) adapt to ligand binding, altering their protonation states to accommodate binding of telaprevir. We suggest that binding of other α-ketoamide inhibitors can lead to the same protonation state changes of the active site histidine residues. Thus, by studying the role of active site protonation changes induced by inhibitors we provide crucial insights to help guide rational drug design, allowing precise tailoring of inhibitors to manipulate the electrostatic environment of SARS-CoV-2 M^pro^.

## INTRODUCTION

The number of confirmed COVID-19 cases worldwide is relentlessly marching towards one hundred million, while the number of deaths is approaching a grim milestone of two million. Sadly, this deadly disease caused by the novel coronavirus SARS-CoV-2 (Severe Acute Respiratory Syndrome Coronavirus 2)^1-4^ has become one of the leading causes of death on the planet in 2020, according to the World Health Organization (www.who.int). Although several vaccines have been developed^5-7^ to slow the spread of SARS-CoV-2, the need for therapeutic intervention options, including small-molecule drugs that inhibit essential steps in the viral replication cycle, cannot be overstated.^8-12^ Small-molecule drugs have shown tremendous success in treating people infected with HIV,^13,14^ hepatitis C^15,16,^ and influenza^17,18^ viruses, and an RNA polymerase inhibitor remdesivir has been recently approved for treatment of COVID-19 by the US Food and Drug Administration.^19^

SARS-CoV-2, a single-stranded, positive-sense RNA virus with a genome comprised of ∼ 30k nucleotides, belongs to the β-coronavirus genus of the *Coronaviridae* family.^20^ A vital step in the viral replication cycle is the cleavage of two polyproteins, pp1a and pp1ab, encoded in the viral replicase gene into individual functional viral proteins.^20,21^ Each polyprotein is mainly processed, or hydrolyzed, by a chymotrypsin-like protease, 3CL M^pro^ or main protease, that belongs to the class of cysteine protease enzymes.^22,23^ The functional main protease (hereafter referred to as M^pro^) is essential for SARS-CoV-2 proliferation as the production of infectious virions depends entirely on the enzymatic activity of M^pro^. Hence, SARS-CoV-2 M^pro^ is undeniably a crucial target for designing specific small-molecule protease inhibitors^24-29^and for potential repurposing of known clinical drugs.^30-35^ Though no clinical drugs are available for use against SARS-CoV-2 M^pro^, several protease inhibitors have been designed to inhibit the very closely related SARS-CoV M^pro36-39^ that shares 96% of amino acid sequence identity with the SARS-CoV-2 enzyme and has a similar catalytic efficiency and almost identical three-dimensional structure. ^25,27,40,41^

Two identical protomers of SARS-CoV-2 M^pro^, each with a molecular mass of ∼34 kDa, create the catalytically active homodimeric enzyme through non-covalent interactions (Fig 1A). Each protomer consists of three structural/functional domains – catalytic domains I (residues 8-101) and II (residues 102-184), and the β-helical domain III (residues 201-303) crucial for protein dimerization (Fig. 1B). Previous studies have shown that the monomeric enzyme is catalytically inactive, as was demonstrated for SARS-CoV M^pro^.^42,43^ The active site cavity is a shallow cleft located on the protein surface between domains I and II. There are six substrate binding subsites, named S1’ through S5, that can bind either substrate residues or chemical groups of inhibitors in positions P1’ through P5. Peptide bond cleavage is carried out at the base of the well-defined subsite S1, where the non-canonical catalytic dyad composed of Cys145 and His41 is located. Catalysis is believed to be assisted by a water molecule positioned at the protein interior side of subsite S2 and hydrogen bonded to the catalytic His41, His164, and Asp187.^25,27,34,40^ Scissile peptide bond cleavage begins through a nucleophilic attack by the Cys145 thiolate on the substrate carbonyl carbon. The negatively charged oxygen of the resultant hemithioketal intermediate is stabilized by a canonical oxyanion hole formed by the main chain amide NH groups of Gly143, Ser144, and Cys145.^44^ Interestingly, subsites S2 and S4 need to be carved out by the substrate or inhibitor groups P2 and P4, respectively, that push protein residues away from their positions in the ligand-free enzyme.^45^ Conversely, subsites S1’, S3, and S5 are fully exposed to the bulk solvent.

**Figure 1.**
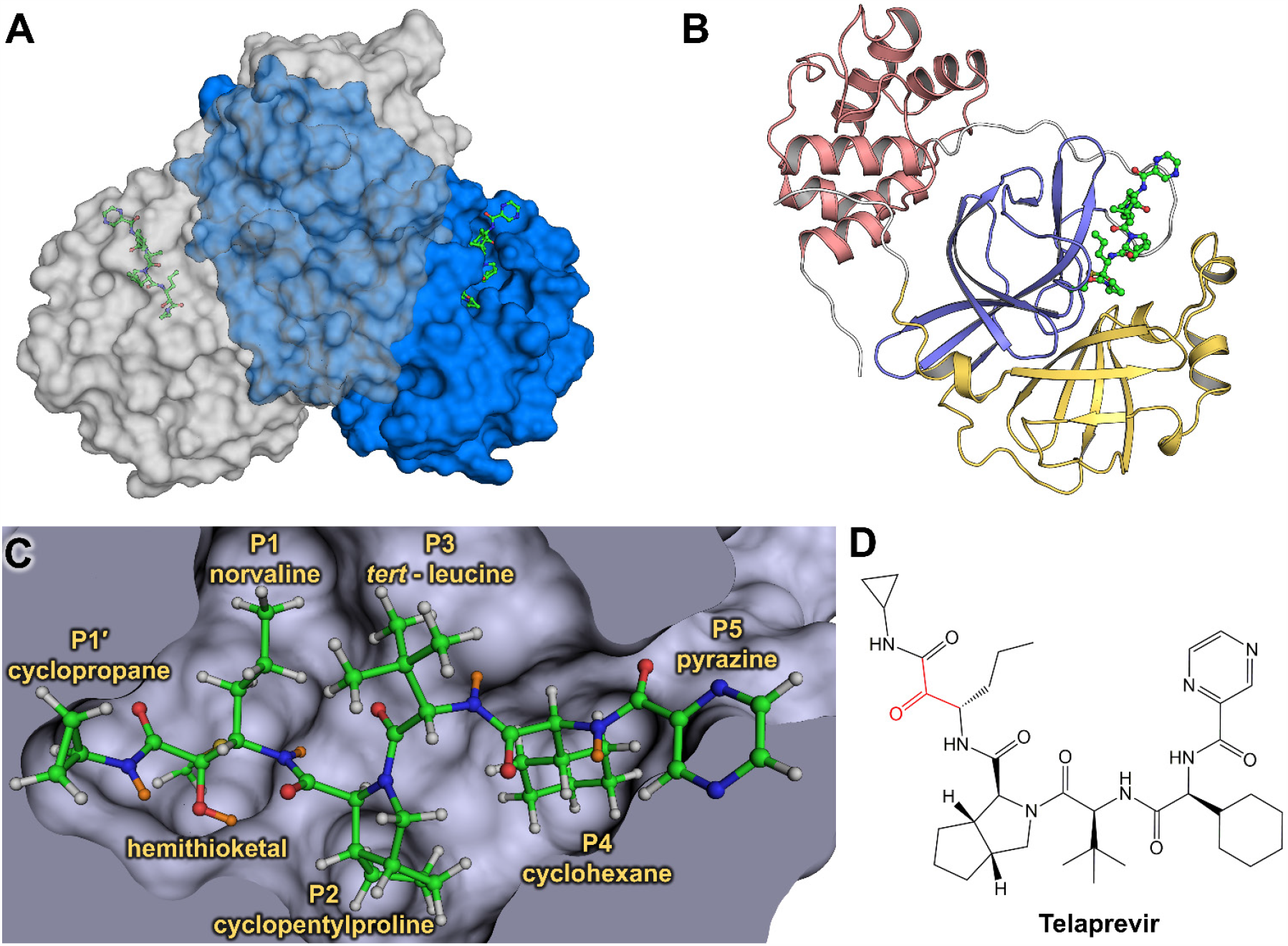
Joint X-ray/neutron structure of SARS-CoV-2 3CL M^pro^ and binding of hepatitis C clinical protease inhibitor telaprevir. (**A**) The catalytically active dimer is shown in surface representation, with telaprevir shown in ball-and-stick representation. (**B**) One enzyme protomer is shown in cartoon representation colored according to the domain structure – domain I is yellow, domain II is blue, and domain III is dark pink. (**C**) Active site cavity with the covalently bound telaprevir. H atoms are shown in gray, whereas D atoms are colored orange. (**D**) A chemical diagram of telaprevir with the ketone warhead shown in red.

Our recent neutron crystallographic study of the ligand-free SARS-CoV-2 M^pro^ provided direct visualization of hydrogen (H) atom locations and hydrogen bonding interactions throughout the enzyme.^46^ The catalytic dyad was observed in a zwitterionic form in the enzyme without substrate or inhibitor. The zwitterion is comprised of a deprotonated, negatively charged Cys145 thiolate and a doubly protonated, positively charged His41 imidazolium situated ∼ 3.8 Å apart from each other. Neutrons are unique probes of light atoms such as H as they are scattered by atomic nuclei instead of electron clouds, which interact with X-rays. Therefore, neutron crystallography can be used to accurately determine the positions of H and its heavier isotope deuterium (D) atoms in biomacromolecules.^47,48^ Deuterium has a potent neutron scattering length of 6.671 fm (https://www.ncnr.nist.gov/resources/n-lengths/), allowing its detection in protein structures even at moderate resolutions of 2.5-2.6 Å.^49-51^ Also, cold neutrons used in neutron crystallographic experiments with wavelengths in the range of 2-5 Å cause no radiation damage to biological samples; therefore, neutron diffraction data can be collected at near-physiological (room) temperature avoiding possible artifacts produced during cryo-cooling and cryo-protection of protein crystals necessary for the mitigation of radiation damage in X-ray cryo-crystallography.^52^

Structure-guided and computer-assisted drug design approaches require a detailed atomic picture of a target biomacromolecule, which is generally obtained using X-ray cryo-crystallography. However, half of the atoms, namely H atoms, are often overlooked in X-ray structures because they usually cannot be located with confidence unless X-ray data extend to sub-Å resolution. Even then, the most interesting and functionally relevant H atoms are typically invisible in electron density maps at ultra-high resolutions because they are mobile and electron-poor, often participating in highly polarized chemical bonds.^53-55^ Detailed structures of the ligand-free and ligand-bound drug targets are essential to steer drug design efforts in the right direction. Knowledge of where H atoms relocate due to inhibitor binding can provide critical information on how protonation states and thus electric charges are modulated in the protein active site cavity, improving rational drug design. For example, molecular dynamics simulations have demonstrated that protonation states in the SARS-CoV-2 M^pro^ active site cavity can be altered when an inhibitor binds,^56^ but experimental evidence has been lacking.

We recently obtained room-temperature X-ray structures of the SARS-CoV-2 M^pro^ in complex with hepatitis C clinical protease inhibitors boceprevir, narlaprevir, and telaprevir, establishing their mode of binding and mechanism of action.^57^ Using *in vitro* enzyme inhibition assays, we determined that these clinical drugs designed against hepatitis C NS3/4A protease inhibited the SARS-CoV-2 M^pro^ activity in the micromolar range. Here, we report a room-temperature neutron structure of the SARS-CoV-2 M^pro^ in complex with telaprevir (M^pro^-Telaprevir) determined at 2.4 Å resolution and refined jointly with a room-temperature X-ray dataset collected from the same crystal to 2.0 Å resolution, thereby improving the accuracy of the experimental model (Table S1).^58^ We chose this complex because it produces crystals of morphology and size amenable for neutron diffraction (Figure S1). Telaprevir represents a promising class of covalent protease inhibitors called α-ketoamides, while also possessing chemical groups in positions P1’ through P5 (Fig. 1C,D).^27^ Locations of D atoms in M^pro^ were accurately determined and compared to those observed in our previous neutron structure^46^ of the ligand-free enzyme, which was crystallized at the same pD value. We discovered that protonation states of key histidine residues (His41, His163, and His164) in the SARS-CoV-2 M^pro^ active site cavity are altered after telaprevir binds, revealing different electric charges between the ligand-free and ligand-bound states of the enzyme. Remarkably, the net electric charge of +1 in the active site cavity is conserved between the two structures. These results are crucial for continued efforts in structure-guided and computational drug design, emphasizing the essential nature of a full atomic-level visualization of protein structure, function, and inhibition.

## RESULTS

### Protonation states in the catalytic site

The reactive carbonyl warhead of the telaprevir β-ketoamide group is attacked by the Cys145 thiolate nucleophile to generate a hemithioketal. Like other β-ketoamide inhibitors,^27,34,57^ the nucleophilic attack is stereospecific, generating only the *S*-enantiomer, in which the newly formed hydroxyl group of the hemithioketal faces towards His41 (Fig. 2A). The distance between the hemithioketal oxygen and His41 Nε2 is 2.6 Å, implicating a strong hydrogen bond. In our neutron structure of the ligand-free SARS-CoV-2 M^pro^ we observed that the Cys145-His41 catalytic dyad adopts a charge-separated state.^46^ Hence, one might suggest that the hemithioketal-His41 pair could maintain charge separation, keeping the positive charge on His41 and having the deprotonated negatively charged hydroxyl on hemithioketal in M^pro^-Telaprevir. Instead, the nuclear density clearly shows that the hemithioketal hydroxyl is protonated (Fig. 2B), whereas His41 is neutral. A reasonable explanation of this observation is that the hemithioketal oxygen is protonated by His41 through a direct proton transfer either in concert with the Cys145 nucleophilic attack on the carbonyl warhead of telaprevir or following the hemithioketal S-C bond formation. Interestingly, the hydroxyl D atom is not positioned on the straight line connecting the hemithioketal oxygen and His41 Nε2 atoms, but instead the O-D…Nε2 angle is 129° indicating that this hydrogen bond may not be strong (Fig 2C).

**Figure 2.**
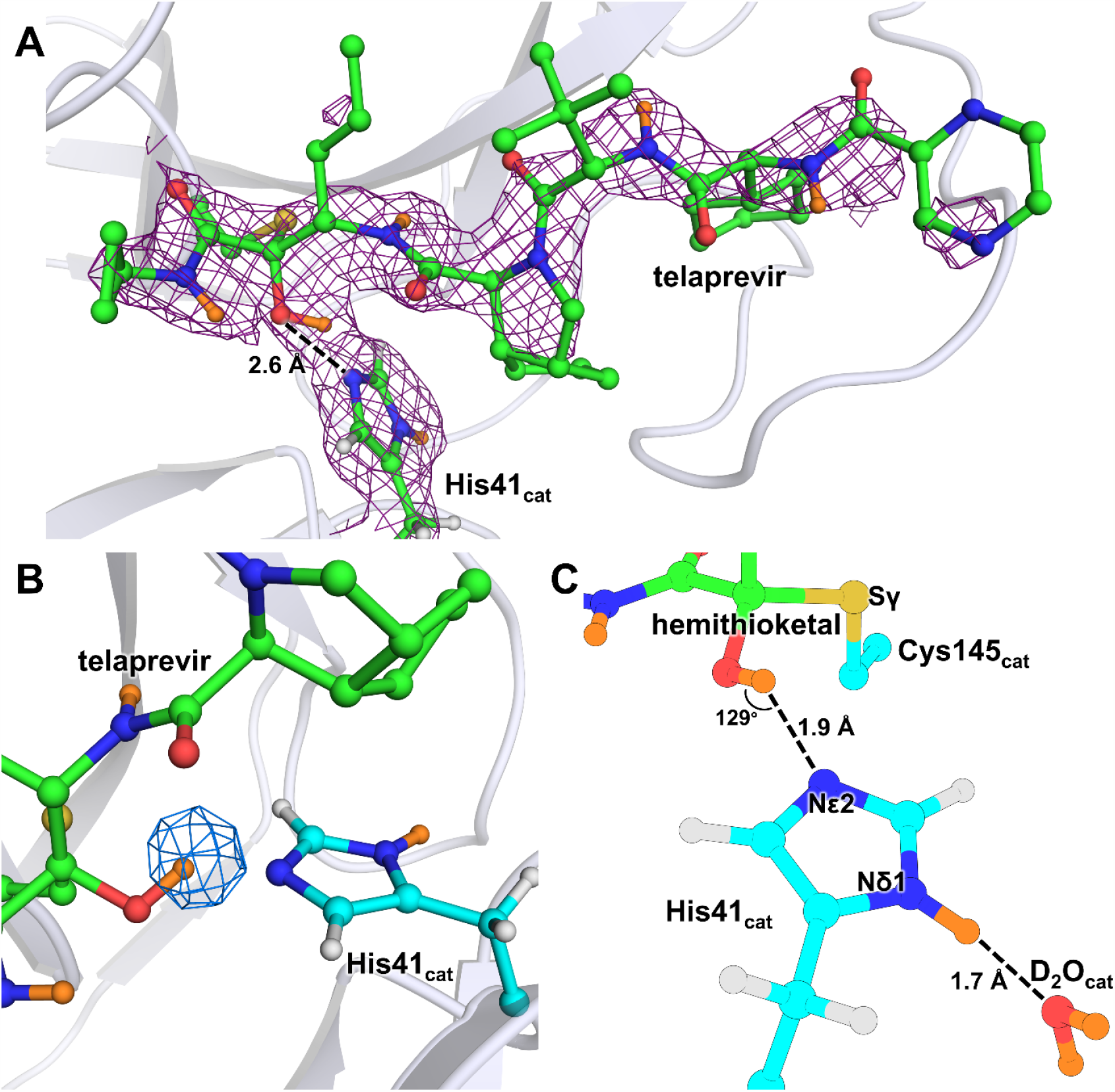
The SARS-CoV-2 M^pro^ catalytic dyad forms a hemithioketal with telaprevir that possesses a protonated hydroxyl. (**A**) The 2FO-FC nuclear density map of telaprevir and the catalytic His41 contoured at a level of 1.0 σ(violet mesh) in the SARS-CoV-2 M^pro^ active site cavity. (**B**) The omit nuclear density map contoured at 3.5 σ level (blue mesh) clearly indicates the hydroxyl of the hemithioketal is protonated. (**C**) The hydrogen bonding interactions involving His41 are shown in detail. H atoms on telaprevir are omitted for clarity, whereas D atoms are colored orange.

In M^pro^-Telaprevir, the His41 imidazole Nδ1 forms a 2.7 Å hydrogen bond with the catalytic water molecule (Nδ1-D…OD_2_O distance of 1.7 Å) (Fig. 2C), which is nearly identical to that observed in the ligand-free M^pro^. The catalytic water is also hydrogen bonded with His164 and Asp187, but it is 3.5 Å away from the main chain ND of His41 compared to 3.2 Å in the ligand-free M^pro^. Thus, the latter hydrogen bond is not present when telaprevir binds. Moreover, in the inhibitor complex, His164 is neutral, having lost the Nε2 D atom that was observed in ligand-free M^pro^. This leads to an increase in Nε2_His164_…Oγ1_Thr175_ distance of 0.2 Å. Hence, the hydrogen bond between His164 and Thr175 is effectively abolished upon telaprevir binding. The hydroxyl of Thr175 remains rotated towards and donates its D in a hydrogen bond with the main chain carbonyl of Asp176. Consequently, in our M^pro^-Telaprevir structure, the catalytic water molecule is no longer surrounded by positively charged histidine side chains, significantly altering the electrostatics around the catalytic site.

### Protonation states in subsite S1

Important ionizable residues of subsite S1 are shown in Figure 3A. Subsite S1 is bordered by residues 140-144 spanning the oxyanion hole, His163, Glu166, and the N-terminal Ser1’ of the other enzyme protomer. Subsite S1 is selective for Gln at substrate position P1. Inhibitors possessing sub-μM affinity to M^pro^ typically mimic Gln by introducing a lactam functionality as their P1 substituent in order to engage His163 in a hydrogen bond.^39^ Telaprevir features a hydrophobic norvaline substituent in P1 position that cannot make hydrogen bonds in subsite S1. Compared to the ligand-free structure, inhibitor binding leads to the recruitment of a water molecule into subsite S1. This water molecule is positioned between the telaprevir’s P1 norvaline and His163 and is hydrogen bonded with the His163 imidazole. Both the electron and nuclear densities for this water molecule are weak, indicating its mobility. Despite the unfavorable norvaline P1 moiety, His163’s protonation state has changed such that the imidazole is doubly protonated and positively charged in M^pro^-Telaprevir compared to that in the ligand-free enzyme, where the Nδ1 not facing the S1 subsite was found to be protonated.^46^ This is significant because His163 protonation explains the apparent ability of the P1 lactam carbonyl to act as a hydrogen bond acceptor with the His163 imidazole observed in several previous X-ray structures of other M^pro^ inhibitors.^26,27,34^ Notably, superposition of M^pro^-Telaprevir with the X-ray structure of inhibitor 13b complex (PDB ID 6Y2F)^27^ reveals that the water molecule position observed in our neutron structure mimics the carbonyl oxygen of inhibitor 13b (Figure S2).

**Figure 3.**
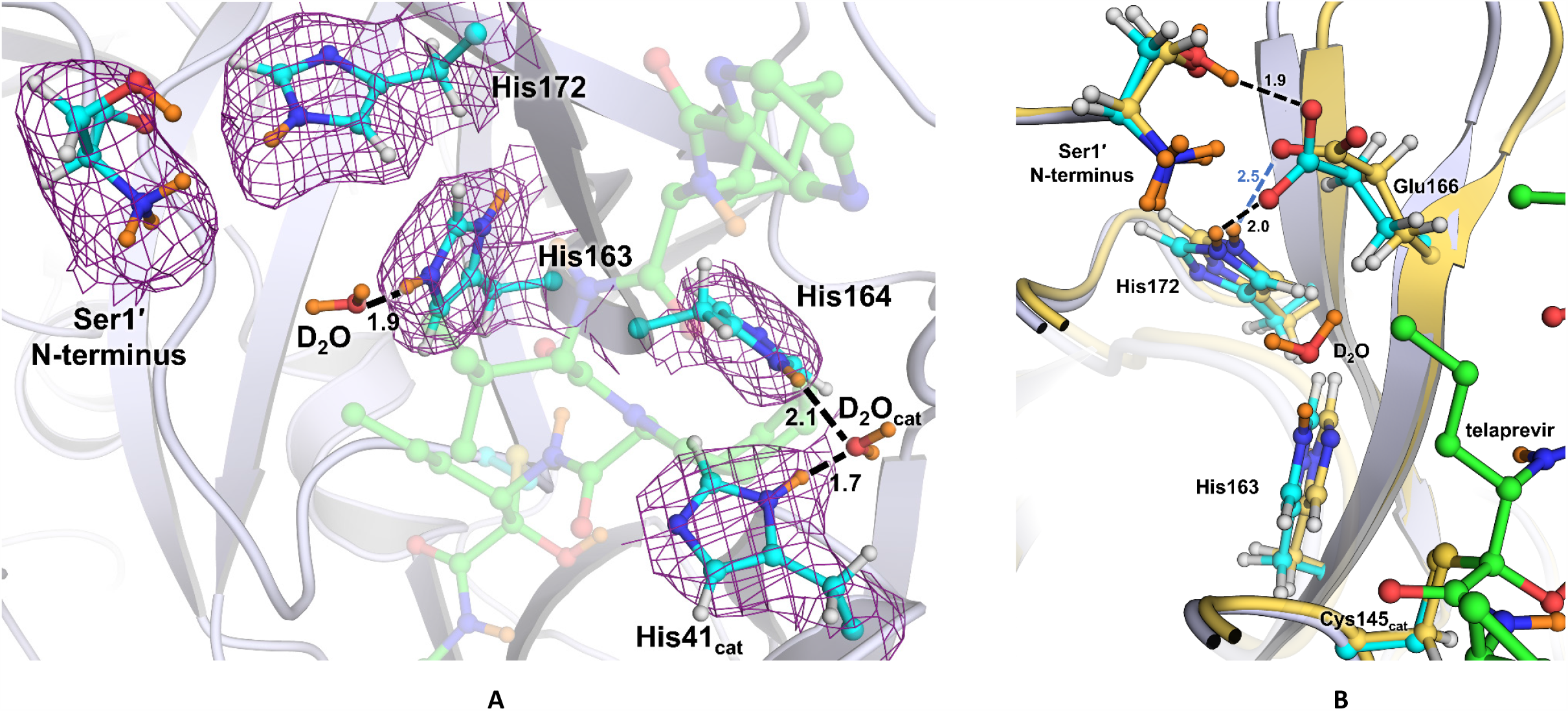
(**A**) Ionizable residues of the active site and S1 subsite of the M^pro^-telaprevir complex. A 2FO-FC nuclear density map of Ser1’ N-terminus and histidines 41,163, 164, and 172 is shown contoured at 1.0 σ level (violet mesh). All distances are shown in Ångstroms with telaprevir being transparent for clarity. (**B**) Superposition of M^pro^-telaprevir and apo-M^pro^ (PDB ID 7JUN) neutron structures showing S1 subsite. Distances are in Å.

Similar to the ligand-free enzyme, the N-terminal amine is found to be protonated and positively charged (ND_3_+) in M^pro^-Telaprevir, where D atoms form a hydrogen bond with the Phe140 main chain carbonyl, the Glu166 carboxylate, and a D_2_O molecule. However, the Glu166 carboxylate side chain is rotated by almost 90° relative to its conformation in the ligand-free enzyme, moving it 0.5 Å towards His172 (Figure 3B). Thus, the Oε1_Glu166…_Nε2_His172_ distance is reduced from 3.3 Å to 2.8 Å, shortening the D…O distance from 2.5 Å to 2.0 Å upon telaprevir binding and considerably strengthening this hydrogen bond. Furthermore, Oε_Glu166_ in the telaprevir complex is 1.2 Å closer to Nε2_His163_ than in ligand-free M^pro^, in agreement with His163 becoming protonated in the M^pro^-Telaprevir inhibitor complex.

## DISCUSSION

Crystallographic structures are essential for making well-educated predictions when designing inhibitors utilizing computer-assisted and structure-guided drug design approaches. It is crucial to determine the locations of heavy atoms and the lighter hydrogen atoms that govern the protonation states of amino acid residues and inhibitor substituents.^55^ The presence or absence of hydrogen atoms on ionizable chemical groups, *e.g.* imidazole of histidine, amine of lysine or N-terminus, or carboxylate of aspartate or glutamate, governs their electric charges and thereby regulates the electrostatics in the vicinity of these groups. Detecting hydrogen atoms in a drug target and a bound ligand is undoubtedly essential to establish protonation states and hydrogen bonding interactions and decipher how protonation states may change due to ligand binding. The potent scattering of neutrons from hydrogen and its isotope deuterium enables neutron crystallography to directly and accurately map hydrogen atoms in biological macromolecules and bound ligands.^46,49-51,59,60^ In this work, we have successfully determined a room-temperature neutron structure of SARS-CoV-2 M^pro^ in complex with the hepatitis C clinical protease inhibitor, telaprevir, from a 0.5 mm^3^ crystal using a partially deuterated enzyme. This structure has permitted us to perform a detailed analysis of D atom positions and compare the protonation states in the active site cavity between the ligand-bound and ligand-free states of the M^pro^ with broad implications for drug design.

In the SARS-CoV-2 M^pro^-Telaprevir neutron structure, we observed protonation of the hemithioketal hydroxyl, likely occurring through a proton transfer from His41 that was found doubly protonated in our previous neutron structure of the ligand-free enzyme. The hemithioketal hydroxyl makes a short, but probably weak, hydrogen bond with the neutral His41 in M^pro^-Telaprevir that would be required to enhance inhibitor binding affinity. Stronger hydrogen bonds would possibly form when the hemithioketal or hemithioacetal oxygen from a warhead carbonyl is directed into the oxyanion hole upon covalent attack on the inhibitor, as was observed in the structures of some M^pro^ inhibitors.^26,34^ The neutral His41 position is stabilized by an additional hydrogen bond with the catalytic water molecule, whose position is held by hydrogen bonds with Asp187 and His164. His164, positioned deeper in the protein interior and observed as doubly protonated in the ligand-free enzyme, has lost its Nε2 D atom resulting in the imidazole ring moving slightly away from its hydrogen bond partner Thr175 upon telaprevir binding. Intriguingly, His163 located in the subsite S1 has gained a proton on its Nε2 atom that faces the bulk solvent in the ligand-free enzyme and P1 norvaline in the telaprevir-bound complex. Moreover, His163 protonation, evidently driven by inhibitor binding, induces a conformational change of the Glu166 side chain that brings it closer to the His163 imidazolium while significantly shortening its hydrogen bond with His172 and retaining the hydrogen bond with the N-terminal ammonium of the second protomer. This observation agrees with His163 becoming positively charged, attracting the negatively charged Glu166. The establishment of hydrogen bond interactions between His163 and the inhibitor lactam ring in several X-ray structures of other inhibitor complexes implies that His163 is protonated in these complexes as Well.^25-27,34,61^

We have observed that the binding of a covalent inhibitor can modulate the protonation states of histidine residues in the active site cavity of SARS-CoV-2 M^pro^. His41, His163, and His164 have different protonation states relative to those in the ligand-free enzyme (Table 1). His41 and His164 lose hydrogens to become uncharged and neutral, whereas His163 gains a proton to become positively charged when telaprevir binds. Cys145 is also uncharged as it is covalently bonded to telaprevir. Nevertheless, the overall electric charge of the active site cavity of +1 is maintained upon inhibitor binding, even though the electrostatics of the ligand-binding cavity differ significantly between the ligand-free and ligand-bound enzyme. It is not unreasonable to suggest that the same protonation state changes of the active site histidine residues may occur upon binding of other α-ketoamide inhibitors. Furthermore, we note that the exact protonation states of the ionizable residues we observed in the active site cavities of ligand-free^46^ and inhibitor-bound SARS-CoV-2 M^pro^ have not been predicted by molecular simulations,^56^ emphasizing the critical importance of experimentally determining the locations of H atoms. As a result, the design of covalent inhibitors against SARS-CoV-2 M^pro^ should consider the observed protonation state changes of the ionizable residues in the active site cavity triggered by inhibitor binding.

**Table 1.** Protonation states and the corresponding electric charges of the ionizable residues in SARS-CoV-2 M^pro^ as observed in the neutron structures of the ligand-free and telaprevir-bound enzyme.

| Residue | **M<sup>pro</sup> ligand-free** (PDB ID 7JUN) |  | **M<sup>pro</sup>-Telaprevir** (PDB ID 7LB7) |  |
| --- | --- | --- | --- | --- |
|  | Charge | Species | Charge | Species |
| Cys145<sub>cat</sub> | −1 | Thiolate (S<sup>−</sup>) | 0 | S-C-OD (hemithioketal) |
| His41<sub>cat</sub> | +1 | Nδ1-D, Nε2-D | 0 | Nδ1-D |
| His163 | 0 | Nδ1-D | +1 | Nδ1-D, Nε2-D |
| His164 | +1 | Nδ1-D, Nε2-D | 0 | Nδ1-D |
| His172 | 0 | Nε2-D | 0 | Nε2-D |
| Net charge | **+1** |  | **+1** |  |

## CONCLUSION

Enzymes rely upon the transfer and movement of hydrogen atoms to carry out their function. We have shown that the active site cleft of SARS-CoV-2 M^pro^ contains three histidine residues that can change their protonation states upon an inhibitor binding, revealing how M^pro^ can tune its electrostatics to accommodate inhibitor and substrate binding into the active site cleft. We observed that inhibitor binding results in a cascade of protonation/deprotonation (Table 2) and conformational events that remodel the SARS-CoV-2 M^pro^ active site cavity as follows:

1. Upon hemithioketal formation a proton is transferred from doubly protonated His41 to the hemithioketal hydroxyl to retain the overall neutral charge of the catalytic site.
2. His164 loses a proton from the Nε2 atom, with no charge, and moves away from Thr175 side chain hydroxyl, which in effect eliminates the hydrogen bond between these two residues.
3. His163 obtains a proton to become doubly protonated, recruiting a loosely bound water molecule to Nε2. The positively charged imidazolium cation attracts the negatively charged carboxylate of Glu166 that changes its conformation to move 1.2 Å closer to His163 and concomitantly strengthens its hydrogen bond with His172.
4. His172 remains singly protonated on Nε2, with no charge, because Nδ1 is locked in a hydrogen bond with the main chain NH of Gly138.
5. The overall +1 electric charge of the active site cavity is retained.

By considering how the protonation states and electric charges of the ionizable histidine residues in the active site can be altered by inhibitor binding in structure-assisted and computational drug design, protease inhibitors can be improved to specifically target the M^pro^ enzyme from SARS-CoV-2.

## METHODS

### General Information

Protein purification columns were purchased from Cytiva (Piscataway, New Jersey, USA). Crystallization reagents were purchased from Hampton Research (Aliso Viejo, California, USA). Crystallographic supplies were purchased from MiTeGen (Ithaca, New York, USA) and Vitrocom (Mountain Lakes, New Jersey, USA). Telaprevir was purchased from BioVision Inc. (Milpitas, CA, USA). A detailed protocol for hydrogenated enzyme expression, purification, and crystallization of M^pro^ to grow neutron diffraction quality crystals has been published elsewhere.^62^

### Cloning, expression, and purification of partially deuterated SARS-CoV-2 M^pro^

The codon-optimized sequence of M^pro^ from SARS-CoV-2 was cloned into the pD451-SR vector harboring kanamycin resistance (ATUM, Newark, CA) and transformed into chemically-competent *E. coli* BL21 (DE3) cells. Before producing deuterated M^pro^, a frozen glycerol stock was first revived in H_2_O minimal medium. Unlabeled glucose (0.5% w/v) and kanamycin (100 mg/mL) was used in this and all other media. After initial growth in H_2_O minimal medium, the cells were adapted stepwise to minimal medium containing increasing percentages (50,75, and 100%) of D_2_O. The final D_2_O-adapted preculture was used to inoculate the bioreactor vessel to an initial volume of 2.8 L. Following inoculation, the BioFlo 310 bioreactor controller console (Eppendorf, Enfield, CT) was set to maintain the temperature (30 C) and the dissolved oxygen level (>30%). The pD was kept above 7.3 by the controlled addition of sodium deuteroxide solution in D_2_O (10% w/w). Once the initial glycerol was exhausted, the culture was fed with a solution containing 20% (w/v) unlabeled glucose and 0.2% (w/v) MgSO_4_. At an OD600 of ∼8.8, isopropyl β-d-1-thiogalactopyranoside (IPTG) was added to a final concentration of 0.5 mM in order to induce protein expression. The cells were collected ∼13.5 h later by centrifugation at 6,000g for 40 min. After removing the supernatant, the wet cell paste was harvested and stored at −80 C until further use. The protein was purified according to the published procedure.^45^ Upstream of the M^pro^ N-terminus codes for a maltose binding protein (MBP) followed by the protease autoprocessing site SAVLQ↓SGFRK (arrow indicates the autocleavage site) which corresponds to the cleavage position between NSP4 and NSP5 in the viral polyprotein. Downstream to the M^pro^ C-terminus codes for the human rhinovirus 3C (HRV-3C) protease cleavage site (SGVTFQ↓GP), which is connected to a 6xHis tag. The N-terminal flanking sequence is autoprocessed *in vivo* during the expression, whereas the C-terminal flanking sequence is removed upon *in vitro* treatment with HRV-3C protease (Millipore Sigma, St. Louis, MO).

### Crystallization

Initial protein crystallization conditions were discovered by screening conducted at the Hauptman-Woodward Medical Research Institute (HWI).^63^ Crystal aggregates were reproduced using the sitting drop vapor diffusion method using 25% PEG3350, 0.1 M Bis-Tris Ph 6.5 in 20μL drops with 1:1 ratio of the protein:well solution. Aggregates were transformed into microseeds using Hampton Research Seed Beads(tm). For partially deuterated M^pro^-Telaprevir co-crystallization, freshly purified deuterated M^pro^ in 20 mM Tris, 150 mM NaCl, 1 mM TCEP, pH 8.0 was concentrated to ∼10.3 mg/mL and mixed with telaprevir, from 60 mM stocks in 100% DMSO, at a 1:5 molar ratio. After room-temperature incubation for 30 minutes, precipitation in the sample was removed via centrifugation and filtration through a 0.2-micron centrifugal filter. Crystallization was achieved in a Hampton 9-well plate and sandwich box set-up with 50 µL drops of protein mixed with 18% PEG3350, 0.1 M Bis-Tris pH 6.5 at 1:1 ratio seeded with 0.2 µL of microseeds at a 1:200 dilution. After 28 days incubation at 14°C, the crystal used for neutron diffraction data collection grew to final dimensions of ∼1.5×0.7×.05 mm (∼0.5 mm^3^) (Figure S1). The crystal was mounted in a fused quartz capillary accompanied with 20% PEG3350 prepared with 100% D_2_O and allowed to H/D exchange for two weeks before starting the neutron data collection. The pH in the crystallization drop at the time of crystal mounting was measured by microelectrode to be 6.6, corresponding to a final pD of 7.0 (pD = pH + 0.4); these are identical conditions to those previously used to determine the neutron structure of the ligand free enzyme.^46^

### Neutron diffraction data collection

The M^pro^-Telaprevir crystal was screened for diffraction quality using a broad-bandpass Laue configuration using neutrons from 2.8 to 10 Å at the IMAGINE instrument at the High Flux Isotope Reactor (HFIR) at Oak Ridge National Laboratory.^64-67^ The full neutron diffraction dataset was then collected using the Macromolecular Neutron Diffractometer (MaNDi) instrument at the Spallation Neutron Source (SNS).^67-69^ The crystal was held stationary at room temperature, and diffraction data were collected for 20 hours using all neutrons between 2-4.16 Å. The next 20-hour diffraction image was collected after crystal rotation by Δϕ = 10°. A total of twenty-one data frames were collected in the final neutron dataset. Diffraction data were reduced using the Mantid package, with integration carried out using three-dimensional TOF profile fitting.^70^ Wavelength normalization of the Laue data was performed using the Lauenorm program from the Lauegen suite.^71,72^ The neutron data collection statics are shown in Table S1.

### X-ray diffraction data collection

The room-temperature X-ray diffraction dataset was collected from the same crystal following the neutron data collection on a Rigaku HighFlux HomeLab instrument equipped with a MicroMax-007 HF X-ray generator and Osmic VariMax optics. The diffraction images were collected using an Eiger R 4M hybrid photon counting detector. Diffraction data were integrated using the CrysAlis Pro software suite (Rigaku Inc., The Woodlands, TX). Diffraction data were then reduced and scaled using the Aimless^73^ program from the CCP4 suite^74^; molecular replacement using PDB code 6XQS^57^ was then performed with Molrep from the CCP4 program suite. The protein structure was first refined against the X-ray using *Phenix.refine* from the Phenix^75^ suite of programs to obtain an accurate model for the subsequent X-ray/neutron joint refinement. The X-ray data collection statics are shown in Table S1.

### Joint X-ray/neutron refinement

The joint X-ray/neutron refinement of ligand-free 3CL M^pro^ was performed using *nCNS*^76^, and the structure was manipulated in *Coot*.^77^ After initial rigid-body refinement, several cycles of positional, atomic displacement parameter, and occupancy refinement were performed. The structure was checked for the correctness of side-chain conformations, hydrogen bonding, and orientations of D_2_O water molecules built based on the mFO-DFC difference neutron scattering length density maps. The 2mFO-DFC and mFO-DFC neutron scattering length density maps were then examined to determine the correct orientations of hydroxyl (Ser, Thr, Tyr), thiol (Cys) and ammonium (Lys) groups, and protonation states of the enzyme residues. The protonation states of some disordered side chains could not be obtained directly and remained ambiguous. All water molecules were refined as D_2_O. Initially, water oxygen atoms were positioned according to their electron density peaks and then were shifted slightly in accordance with the neutron scattering length density maps. Because M^pro^ for this study was partially deuterated, all H positions in the protein were modeled as D atoms. In telaprevir only the labile H atoms were modeled as Ds. The occupancies of D atoms were refined individually within the range of −0.56 (pure H) to 1.00 (pure D) because the neutron scattering length of H is –0.56 times that of D. Before depositing the neutron structure to the PDB, a script was run that converts a record for the coordinates of a D atom into two records corresponding to an H and a D partially occupying the same site, both with positive partial occupancies that add up to unity. The percent D at a specific site is calculated according to the following formula: %D= {Occupancy(D) + 0.56}/1.56.

## Data availability

The coordinates and structure factors for the SARS-CoV-2 3CL M^pro^-Telaprevir complex have been deposited in the PDB with the accession code 7LB7. Any other relevant data are available from the corresponding authors upon reasonable request.

## Acknowledgments

This research was supported by the DOE Office of Science through the National Virtual Biotechnology Laboratory (NVBL), a consortium of DOE national laboratories focused on response to COVID-19, with funding provided by the Coronavirus CARES Act. This research used resources at the Spallation Neutron Source and the High Flux Isotope Reactor, which are DOE Office of Science User Facilities operated by the Oak Ridge National Laboratory. The Office of Biological and Environmental Research supported research at ORNL’s Center for Structural Molecular Biology (CSMB), a DOE Office of Science User Facility. This research used resources at the Second Target Station, which is a DOE Office of Science User Facilities Construction Project at Oak Ridge National Laboratory. We thank Dr. Hugh M. O’Neill from ORNL for assistance during expression of the partially deuterated protein

## Author contributions

L.C and A.K. conceived the study. A.K., G.P. and Q.Z. designed and cloned the gene. G.P., K.L.W. and Q.Z. performed expression of the partially deuterated protein. D.W.K. and A.K. crystallized the protein. A.K. and D.W.K. collected the X-ray diffraction data. L.C. collected and reduced the neutron diffraction data. A.K., D.W.K. and L.C. refined the structure. D.W.K., L.C., and A.K. wrote the paper with help from all co-authors.

