## Supplemental Table 1 and Figures S1-2 for "Inhibitor Binding Modulates Protonation States in the Active Site of SARS-CoV-2 Main Protease"

Table S1

| | $M^{pro}$-Telaprevir, RT, PDB ID 7LB7 | |
|---|---|---|
| **Data collection:** | **Neutron** | **X-ray** |
| Beamline/Facility | MaNDi (SNS) | Rigaku HighFlux HomeLab |
| Space group | C2 | |
| Cell dimensions: | | |
| *a*, *b*, *c* (Å) | 110.56, 55.69, 48.81 | |
| α, β, γ (°) | 90, 101.3, 90 | |
| Resolution (Å) | 14.29-2.40 (2.49-2.40)* | 54.21-2.00 (2.05-2.00)* |
| No. reflections measured | 38766 | 80233 |
| No. reflections unique | 9566 (888) | 19852 (1854) |
| $R_{merge}$ | 0.176 (0.336) | 0.082 (0.544) |
| $R_{pim}$ | 0.085 (0.201) | 0.046 (0.307) |
| $CC_{1/2}$ | 0.975 (0.532) | 0.993 (0.670) |
| $I$ / σ$I$ | 11.4 (3.1) | 13.9 (1.9) |
| Completeness (%) | 83.4 (77.4) | 96.3 (93.3) |
| Redundancy | 4.05 (2.68) | 4.2 (4.0) |
| **Refinement:** | **Joint XN** | |
| Resolution (neutron, Å) | 14.29 – 2.40 | |
| Resolution (X-ray, Å) | 27.85 – 2.00 | |
| Data rejection criteria | no observation & \|F\|=0 | |
| Sigma cut-off | 2.5 | |
| No. reflections (neutron) | 9539 | |
| No. reflections (X-ray) | 19261 | |
| $R_{work}$ / $R_{free}$ (neutron) | 0.216 / 0.228 | |
| $R_{work}$ / $R_{free}$ (X-ray) | 0.204 / 0.225 | |
| No. atoms | | |
| Protein, including H and D | 4675 | |
| Telaprevir | 107 | |
| Water | 285 (i.e. 95 $D_2O$ molecules) | |
| *B*-factors | | |
| Protein | 39.8 | |
| Telaprevir | 33.1 | |
| Water | 51.0 | |
| R.M.S. deviations | | |
| Bond lengths (Å) | 0.009 | |
| Bond angles (°) | 1.094 | |

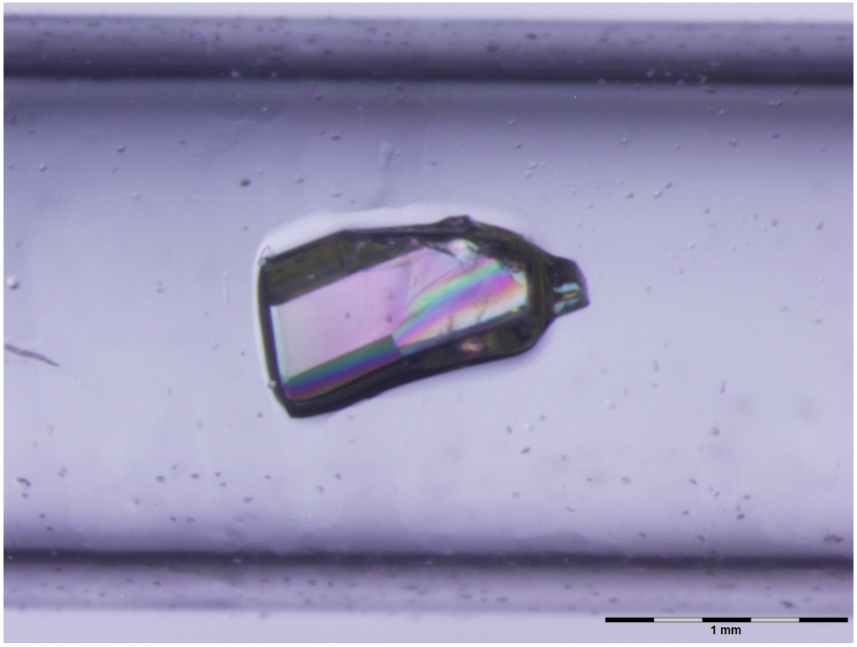


**Figure S1.** The 0.5mm$^3$ crystal of SARS-CoV-2 M$^{pro}$-Telaprevir complex used in the current study.

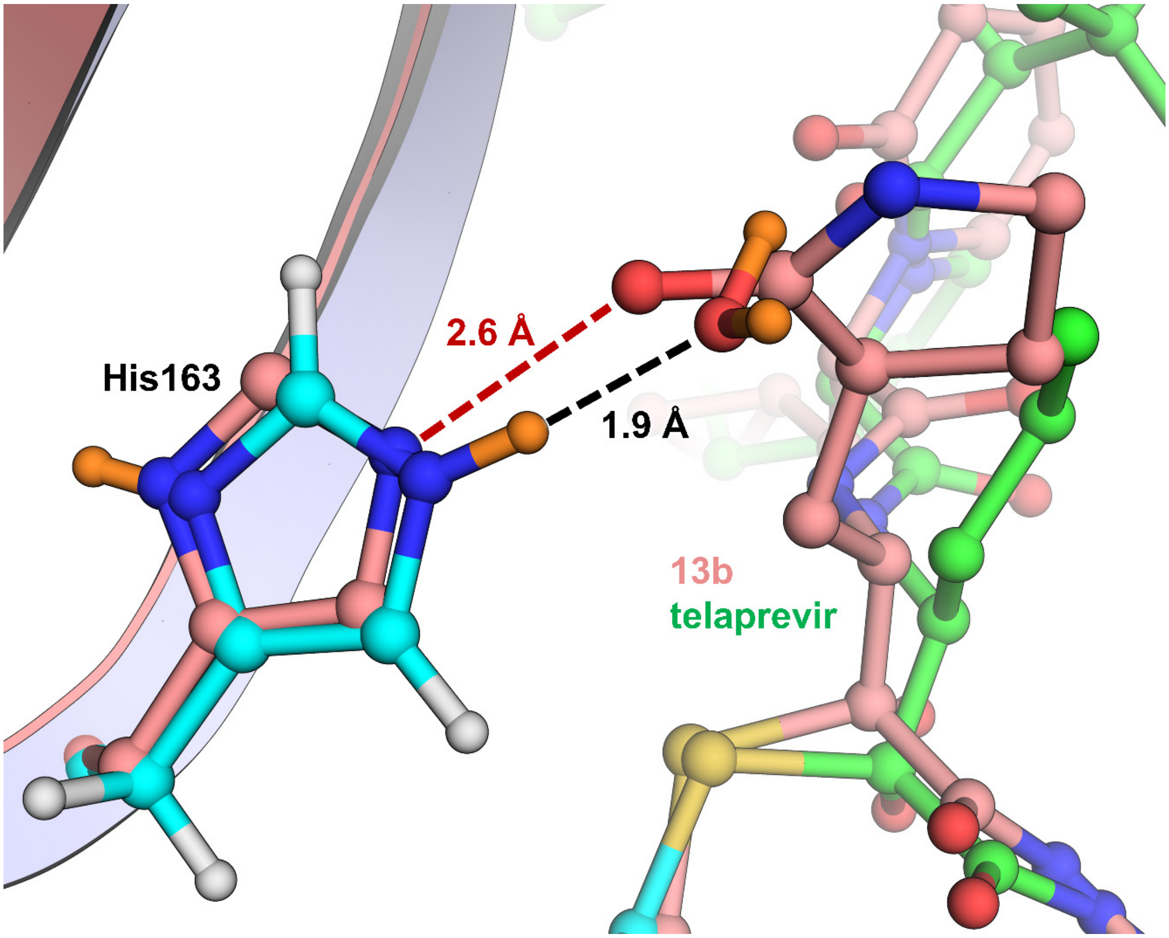


**Figure S2.** Superposition of M$^{pro}$-Telaprevir neutron structure (cyan carbons for protein and green carbons for telaprevir) with the X-ray structure of M$^{pro}$ in complex with inhibitor 13b (PDB ID 6Y2F) showing a $D_2O$ molecule in M$^{pro}$-Telaprevir whose position acts as a template for the carbonyl group of the P1 lactam of 13b.
